## Supplemental tables and figures for "Effect of stimulation time on the expression of human macrophage polarization markers"

### Supporting information

#### S1 Table

| M1 | | | | | | | M2a | | | | | | M2c | | | |
|---|---|---|---|---|---|---|---|---|---|---|---|---|---|---|---|---|
| | CXCL9 | CXCL10 | TNF | IL-1β | IL-12 | IDO1 | | MRC1 | TGM2 | CCL17 | CCL22 | IL-10 | | IL-10 | CD163 | TGFβ |
| US vs. 4h | ** | *** | *** | ** | ** | * | US vs. 4h | *** | *** | * | *** | *** | US vs. 4h | *** | *** | ns |
| US vs. 8h | *** | *** | *** | ** | *** | ** | US vs. 8h | *** | ** | * | *** | *** | US vs. 8h | *** | ** | ns |
| US vs. 12h | ** | ** | *** | *** | *** | *** | US vs. 12h | *** | ** | * | ** | *** | US vs. 12h | ns | *** | ** |
| US vs. 24h | * | ** | *** | * | ns | ns | US vs. 24h | *** | *** | * | *** | *** | US vs. 24h | ** | *** | ns |
| US vs. 48h | ** | *** | *** | ** | *** | ** | US vs. 48h | *** | ** | * | ** | *** | US vs. 48h | *** | *** | *** |
| US vs. 72h | * | ** | *** | * | * | *** | US vs. 72h | *** | ** | * | *** | *** | US vs. 72h | *** | *** | *** |
| 4h vs. 8h | ** | *** | *** | *** | *** | ** | 4h vs. 8h | ns | ns | ns | *** | *** | 4h vs. 8h | ** | ns | ns |
| 4h vs. 12h | ** | *** | *** | * | *** | ** | 4h vs. 12h | * | ns | * | ** | ** | 4h vs. 12h | *** | ** | * |
| 4h vs. 24h | ** | *** | *** | ns | ** | ns | 4h vs. 24h | ns | ns | ** | *** | *** | 4h vs. 24h | *** | ns | ns |
| 4h vs. 48h | ** | *** | *** | ns | ns | ** | 4h vs. 48h | * | ns | ns | ** | *** | 4h vs. 48h | *** | ** | * |
| 4h vs. 72h | ** | *** | *** | *** | ns | *** | 4h vs. 72h | *** | * | ** | *** | *** | 4h vs. 72h | ** | *** | ns |
| 8h vs. 12h | *** | ** | *** | ns | ns | ns | 8h vs. 12h | ns | * | * | * | ns | 8h vs. 12h | *** | ns | ns |
| 8h vs. 24h | *** | *** | *** | ns | ** | ns | 8h vs. 24h | *** | * | * | *** | *** | 8h vs. 24h | ** | ns | ns |
| 8h vs. 48h | ** | *** | *** | ns | *** | ** | 8h vs. 48h | * | ns | ns | ** | *** | 8h vs. 48h | ns | * | ** |
| 8h vs. 72h | *** | *** | *** | * | ** | *** | 8h vs. 72h | *** | *** | * | ** | *** | 8h vs. 72h | ns | * | ns |
| 12h vs. 24h | ns | ns | ** | ns | * | ns | 12h vs. 24h | *** | ns | ns | *** | *** | 12h vs. 24h | ** | ** | ns |
| 12h vs. 48h | ns | ** | ns | ns | ** | ** | 12h vs. 48h | * | ns | * | ** | *** | 12h vs. 48h | *** | *** | * |
| 12h vs. 72h | ** | ** | * | ns | ** | *** | 12h vs. 72h | *** | ** | ns | ** | *** | 12h vs. 72h | * | *** | ns |
| 24h vs. 48h | ns | ** | * | ns | ** | * | 24h vs. 48h | *** | ns | ** | * | *** | 24h vs. 48h | ** | *** | * |
| 24h vs. 72h | * | ** | ** | ns | * | *** | 24h vs. 72h | *** | ** | ns | ** | ns | 24h vs. 72h | ns | * | ns |
| 48h vs. 72h | * | * | *** | * | ns | ns | 48h vs. 72h | *** | ** | ** | ns | *** | 48h vs. 72h | ns | ns | ns |

**S1 Table. Statistical analyses for M1, M2a and M2c markers at mRNA level in polarized macrophages.** Expression of the indicated markers at the time points shown were compared by repeated measures ANOVA, * $p < 0.05$; ** $p < 0.01$; *** $p < 0.001$. ns, not significant; US, unstimulated.

**S2 Table**

| M1 | | | | M2a | | | M2c | |
|---|---|---|---|---|---|---|---|---|
| | CD86 | CD64 | HLA-DR | | CD200R | CD206 | | CD163 |
| US vs. 4h | ns | ns | ns | US vs. 4h | ns | ns | US vs. 4h | ns |
| US vs. 8h | * | ** | ns | US vs. 8h | * | * | US vs. 8h | ns |
| US vs. 12h | * | * | * | US vs. 12h | * | * | US vs. 12h | ns |
| US vs. 24h | ns | * | ns | US vs. 24h | * | ** | US vs. 24h | ns |
| US vs. 48h | ns | * | * | US vs. 48h | * | * | US vs. 48h | ns |
| US vs. 72h | ** | * | ns | US vs. 72h | * | * | US vs. 72h | ns |
| 4h vs. 8h | ** | ** | ns | 4h vs. 8h | ns | ns | 4h vs. 8h | * |
| 4h vs. 12h | * | ns | * | 4h vs. 12h | ns | ns | 4h vs. 12h | * |
| 4h vs. 24h | ns | * | ns | 4h vs. 24h | * | * | 4h vs. 24h | ns |
| 4h vs. 48h | * | * | ns | 4h vs. 48h | ** | * | 4h vs. 48h | ns |
| 4h vs. 72h | ns | * | ns | 4h vs. 72h | * | * | 4h vs. 72h | ns |
| 8h vs. 12h | ns | ns | ns | 8h vs. 12h | * | ns | 8h vs. 12h | ns |
| 8h vs. 24h | ** | ns | * | 8h vs. 24h | * | ** | 8h vs. 24h | ns |
| 8h vs. 48h | * | * | * | 8h vs. 48h | ** | * | 8h vs. 48h | ns |
| 8h vs. 72h | ns | * | ns | 8h vs. 72h | * | ns | 8h vs. 72h | ns |
| 12h vs. 24h | * | ns | * | 12h vs. 24h | ns | ** | 12h vs. 24h | ns |
| 12h vs. 48h | * | * | * | 12h vs. 48h | ** | * | 12h vs. 48h | ns |
| 12h vs. 72h | ns | * | * | 12h vs. 72h | * | ns | 12h vs. 72h | ns |
| 24h vs. 48h | * | ns | ns | 24h vs. 48h | *** | ns | 24h vs. 48h | ns |
| 24h vs. 72h | * | ns | ns | 24h vs. 72h | ns | ns | 24h vs. 72h | ns |
| 48h vs. 72h | ns | ns | ns | 48h vs. 72h | ns | ns | 48h vs. 72h | ns |

**S2 Table. Statistical analyses for M1, M2a and M2c surface markers in polarized macrophages.** Expression of the indicated markers at the time points shown were compared by repeated measures ANOVA, * $p < 0.05$; ** $p < 0.01$; *** $p < 0.001$. ns, not significant; US, unstimulated.

**S3 Table**

| M1 | | | | M2a | | M2c | |
|---|---|---|---|---|---|---|---|
| | IL12p70 | TNF | IL1β | | IL10 | | TGFβ |
| US vs. 4h | *** | *** | ns | US vs. 4h | ** | US vs. 4h | ns |
| US vs. 8h | *** | *** | * | US vs. 8h | ** | US vs. 8h | ns |
| US vs. 12h | *** | *** | * | US vs. 12h | ** | US vs. 12h | - |
| US vs. 24h | *** | *** | * | US vs. 24h | ns | US vs. 24h | ns |
| US vs. 48h | *** | *** | ns | US vs. 48h | *** | US vs. 48h | ns |
| US vs. 72h | *** | ** | ns | US vs. 72h | *** | US vs. 72h | ** |
| 4h vs. 8h | *** | * | ns | 4h vs. 8h | - | 4h vs. 8h | ns |
| 4h vs. 12h | *** | * | * | 4h vs. 12h | - | 4h vs. 12h | ns |
| 4h vs. 24h | *** | ** | * | 4h vs. 24h | ns | 4h vs. 24h | ns |
| 4h vs. 48h | *** | *** | ns | 4h vs. 48h | *** | 4h vs. 48h | ns |
| 4h vs. 72h | *** | *** | ns | 4h vs. 72h | *** | 4h vs. 72h | ** |
| 8h vs. 12h | *** | ns | * | 8h vs. 12h | - | 8h vs. 12h | ns |
| 8h vs. 24h | * | *** | ns | 8h vs. 24h | ns | 8h vs. 24h | ns |
| 8h vs. 48h | ns | *** | ns | 8h vs. 48h | *** | 8h vs. 48h | ns |
| 8h vs. 72h | ns | *** | ns | 8h vs. 72h | *** | 8h vs. 72h | ** |
| 12h vs. 24h | ** | ** | ns | 12h vs. 24h | ns | 12h vs. 24h | ns |
| 12h vs. 48h | * | *** | ns | 12h vs. 48h | *** | 12h vs. 48h | ns |
| 12h vs. 72h | * | *** | * | 12h vs. 72h | *** | 12h vs. 72h | ** |
| 24h vs. 48h | ** | *** | ns | 24h vs. 48h | *** | 24h vs. 48h | ns |
| 24h vs. 72h | * | *** | ns | 24h vs. 72h | *** | 24h vs. 72h | ** |
| 48h vs. 72h | ns | ns | ns | 48h vs. 72h | ns | 48h vs. 72h | ** |

**S3 Table. Statistical analyses for M1, M2a and M2c cytokines in polarized macrophages.** Expression of the indicated markers at the time points shown were compared by repeated measures ANOVA, * $p < 0.05$; ** $p < 0.01$; *** $p < 0.001$. ns, not significant; US, unstimulated.

#### S1 Fig

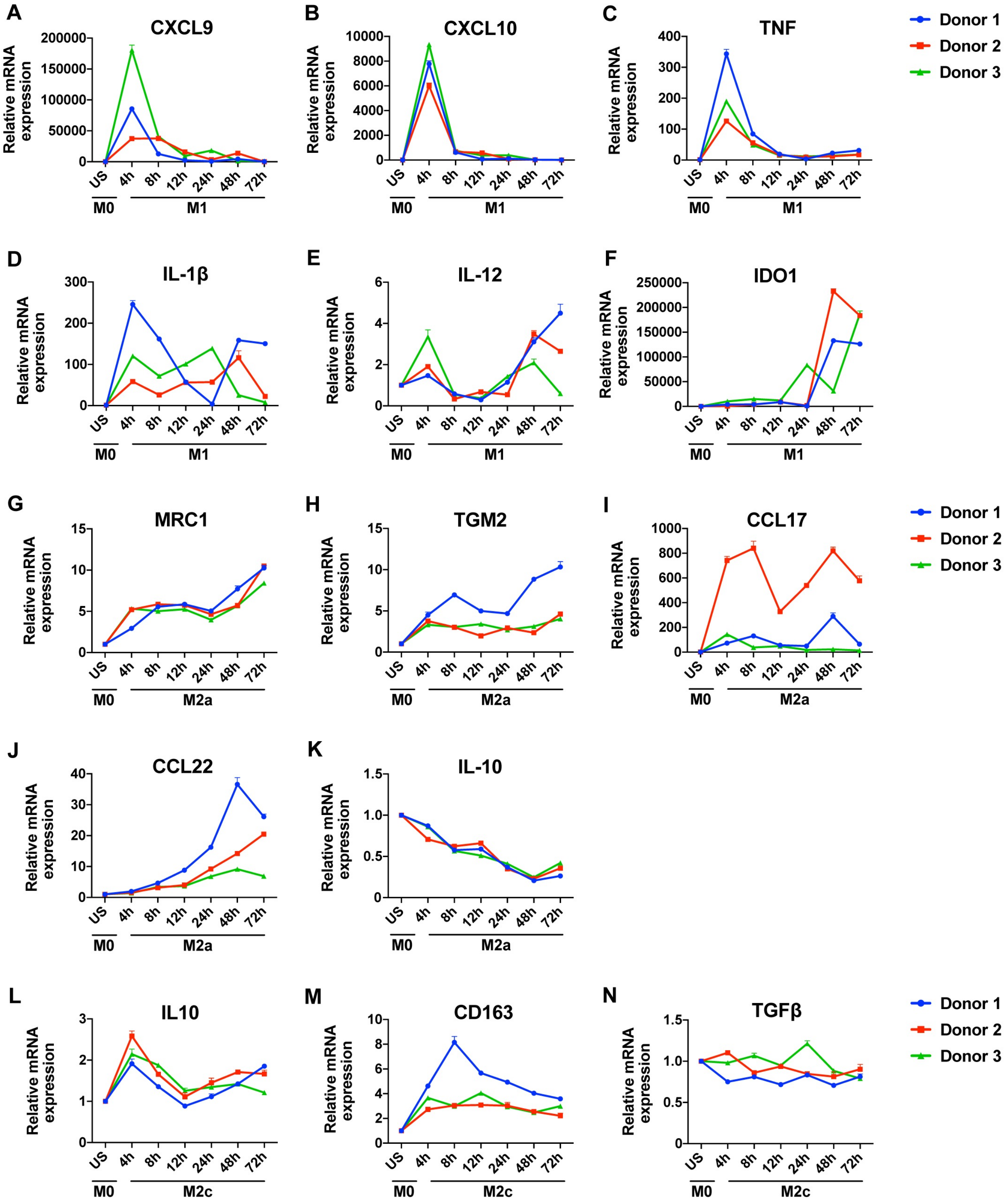


**S1 Fig. Time-dependent changes in the expression of macrophage polarization markers at the mRNA level for each donor.**

#### S2 Fig

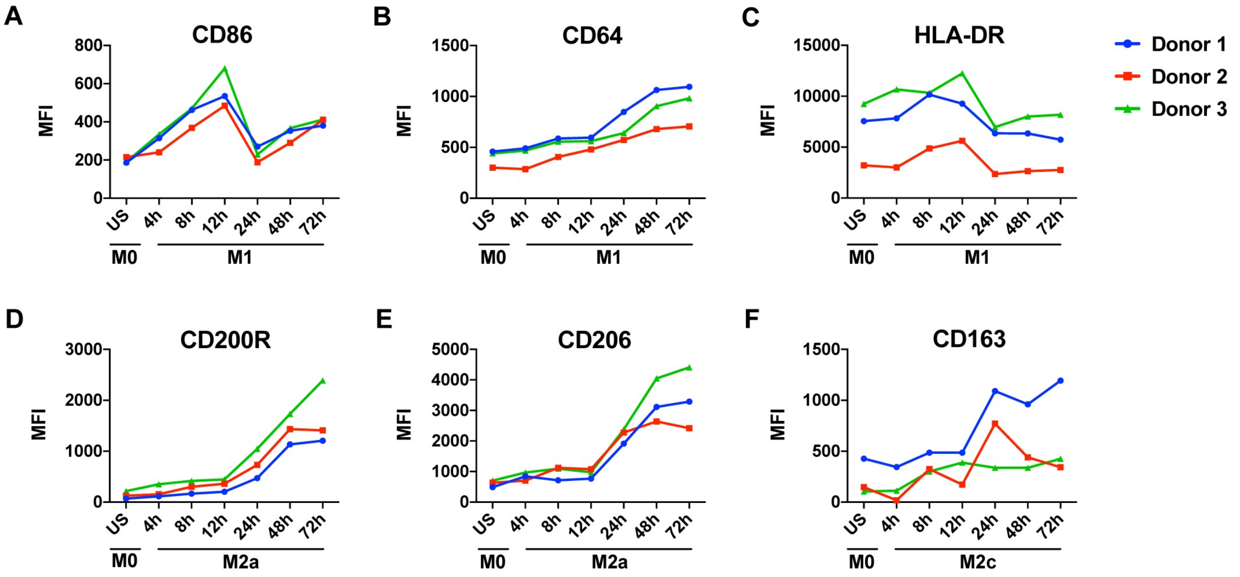


**S2 Fig. Time-dependent changes in the expression of macrophage polarization markers at the protein level for each donor.**

#### S3 Fig

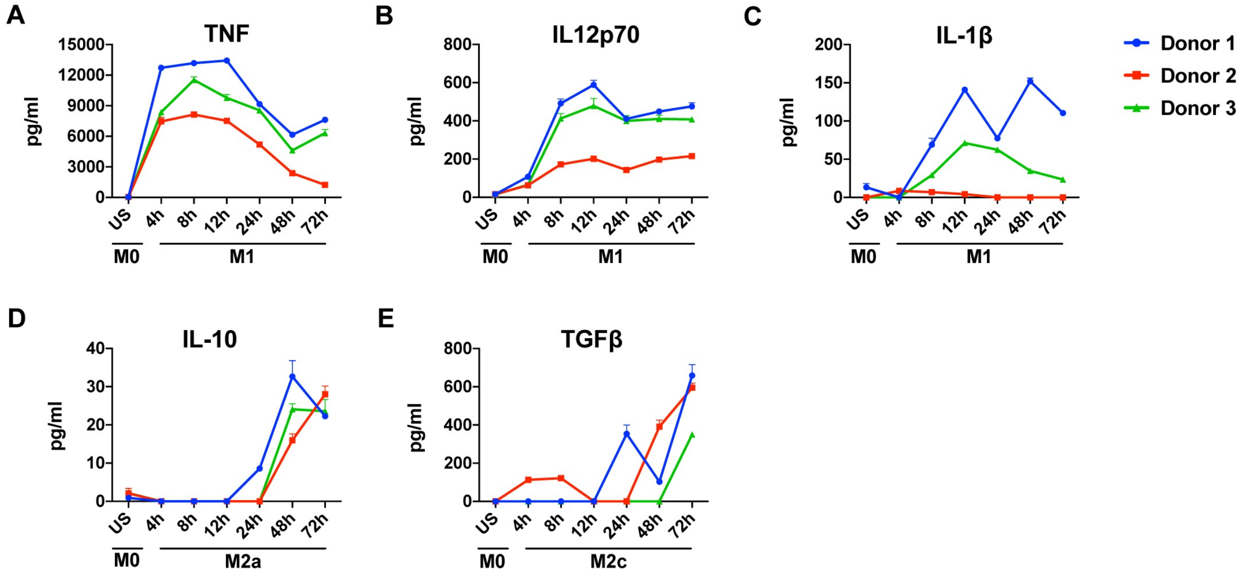


**S3 Fig. Time-dependent changes in cytokine production for each donor.**

#### S4 Fig

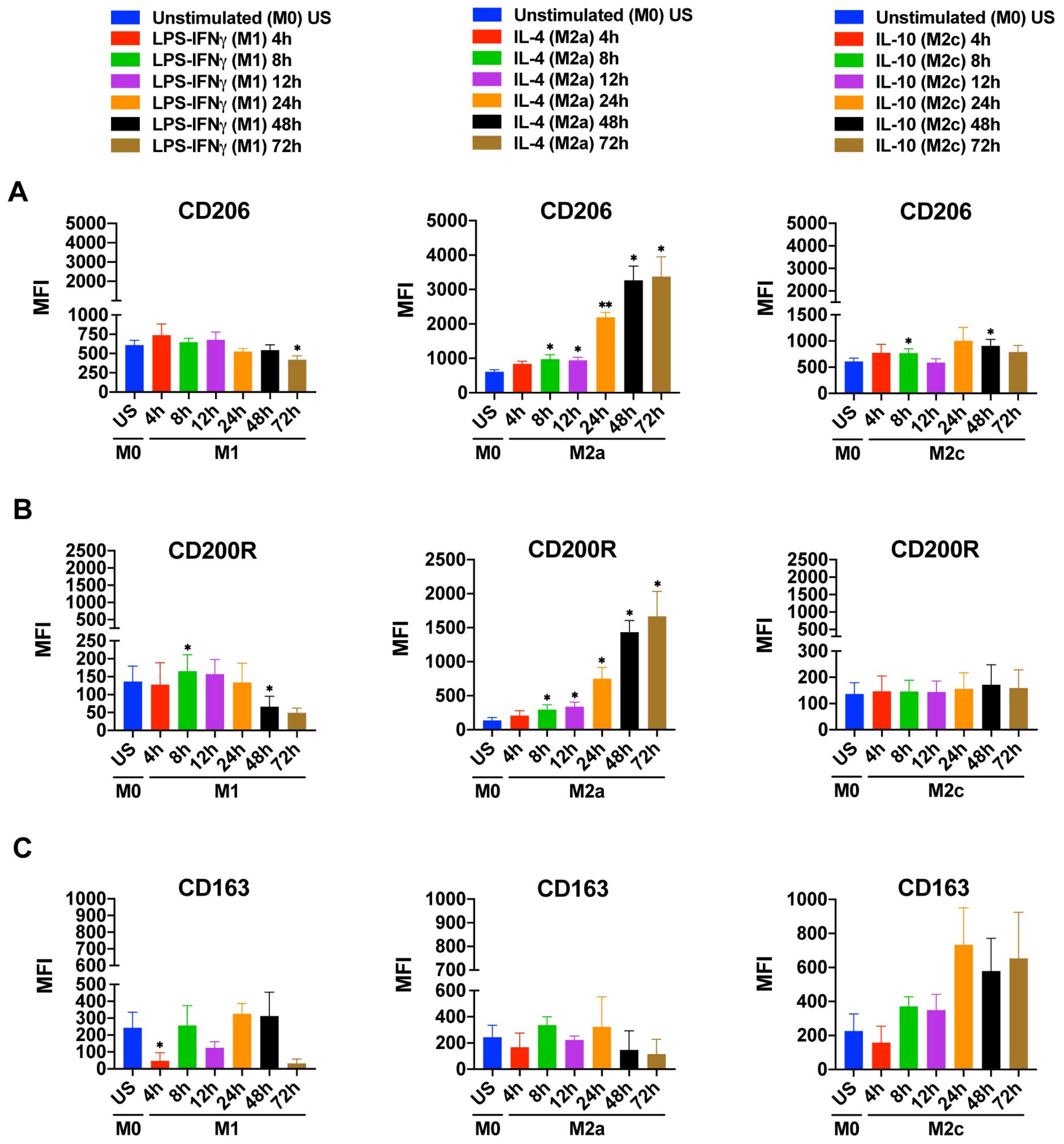


**S4 Fig. Time depended changes in the expression of M2 markers in M1, M2a and M2c macrophages analysed by flow cytometry.** Summary data shown are mean ± SEM of biological replicates of 3 independent donors. Polarized macrophages (M1, M2a, or M2c) at all time points were compared with unstimulated (US) M0 macrophages. Statistical analyses were performed with repeated measures ANOVA, * $p < 0.05$; ** $p < 0.01$; *** $p < 0.001$.
